# Histone Acetyltransferase 1 is Required for DNA Replication Fork Function and Stability

**DOI:** 10.1101/2020.03.17.989434

**Authors:** Paula A. Agudelo Garcia, Callie Lovejoy, Prabakaran Nagarajan, Dongju Park, Liudmila Popova, Michael A. Freitas, Mark R. Parthun

**Affiliations:** Department of Biological Chemistry and Pharmacology, The Ohio State University, Columbus, OH 43210; Department of Cancer Biology and Genetics, The Ohio State University, Columbus, OH 43210

## Abstract

The replisome functions in a dynamic environment that is at the intersection of parental and nascent chromatin. Parental nucleosomes are disrupted in front of the replication fork. The daughter duplexes are packaged with an equal amount of parental and newly synthesized histones in the wake of the replication fork through the action of the replication-coupled chromatin assembly pathway. Histone acetyltransferase 1 (Hat1) is responsible for the cytosolic diacetylation of newly synthesized histone H4 on lysines 5 and 12 that accompanies replication-coupled chromatin assembly. Analysis of the role of Hat1 in replication-coupled chromatin assembly demonstrates that Hat1 also physically associates with chromatin near sites of DNA replication. The association of Hat1 with newly replicated DNA is transient but can be stabilized by replication fork stalling. The association of Hat1 with nascent chromatin may be functionally relevant as loss of Hat1 results in a decrease in replication fork progression and an increase in replication fork stalling. In addition, in the absence of Hat1, stalled replication forks are unstable and newly synthesized DNA becomes susceptible to Mre11-dependent degradation. These results suggest that Hat1 links replication fork function to the proper processing and assembly of newly synthesized histones.

## INTRODUCTION

The central event in the division of a cell is the duplication of its chromosomes. Chromosome duplication requires the proper functioning of two interconnected processes. The first is the replication of the genomic DNA. The second is the duplication of the chromatin structure that governs the correct packaging and architecture of the chromosomes in the nucleus. The successful coordination and completion of these processes is essential to ensure genome stability, maintain correct patterns of gene expression and properly regulate cell proliferation.

DNA replication occurs in a unique and highly dynamic chromatin environment. The replication fork must navigate through the chromatin structure in front of the fork by disrupting nucleosomes in its path. In the wake of the replication fork, the nascent daughter duplexes must be rapidly assembled into nucleosomes. Nascent chromatin on the daughter duplexes is assembled from two distinct pools of histones; parental and newly synthesized.

The parental histones are derived from the nucleosomes disrupted during the passage of the replication fork. These nucleosomes dissociate into stable H3/H4 tetramers and H2A/H2B dimers. Regulation of parental histone recycling, mediated by the histone chaperone Asf1, is critical for proper replication fork function and stability (1). Asf1 functions in conjunction with the MCM2-7 replicative helicase and RPA to remove parental histones H3 and H4 from in front of the replication fork and transfer them to the newly replicated DNA near their original genomic location(2-6). Loss of Asf1 or disruption of Asf1 activity through histone over-expression impedes DNA unwinding and replication fork progression (7-9). Other factors, such as FACT and the POLE3-POLE4 complex are also involved in processing parental histones at the replication fork and may be involved in the association of H2A/H2B dimers with the H3/H4 tetramers(10-12).

To maintain nucleosome density, an equal quantity of newly synthesized histones must be delivered to sites of DNA replication. This is accomplished through the replication-coupled chromatin assembly pathway. The replication-coupled chromatin assembly pathway begins with the processing of new newly synthesized H3 and H4 in the cytoplasm, where there is a large burst of histone synthesis to meet the needs of chromosome duplication(13). The histone methyltransferase SetDB1 is associated with the ribosome and mono-methylates histone H3 lysine 9 co-translationally(14). H3 and H4 then form stable heterodimers in a chaperone-mediated process(15). The H3/H4 dimers are then bound by the Hat1 complex, which consists of the histone acetyltransferase Hat1 and the histone chaperone Rbap46, resulting in the acetylation of H4 on lysines 5 and 12(16-20). The modified H3/H4 dimers are then transferred to the histone chaperone Asf1 through the direct association of Asf1 with the Hat1/Rbap46/H3/H4 complex(15,21). The Asf1/H3/H4 complex can associate with specific importins/karyopherins to facilitate the nuclear import of the H3/H4 dimers(21-26). Once in the nucleus, additional processing occurs on some H3/H4 complexes with the acetylation of H3 lysine 56 by CBP/Rtt109 and additional lysine residues in the H3 NH_2_-terminal tail by Gcn5(21,27-38). The modified H3/H4 complexes are then transferred to the CAF-1 complex (Chromatin Assembly Factor 1), which facilitates the deposition of H3/H4 tetramers near replication forks through a physical interaction with PCNA(39-47).

Several recent studies have shown that replication-coupled chromatin assembly is required for replication fork function. In these studies, the supply of histones to the replication fork was blocked, either by preventing new histone protein synthesis or disrupting histone deposition by depleting CAF-1 or Asf1(39,48-51). There is also evidence that the post-translational modifications on newly synthesized histones can influence replication fork function. Histone H3 lysine 56 acetylation has been shown to positively regulate binding of histones to CAF-1(38). Consistent with this, loss of H3 lysine 56 acetylation and CAF-1 have similar effects on DNA replication in *S. cerevisiae*(49). Histone deacetylases, HDAC1 and HDAC2, which have been proposed to deacetylate newly synthesized histones following their assembly into chromatin, have been shown to be important for replication fork function and for the stabilization of stalled replication forks in conjunction with the WRN helicase(52-56).

Recent evidence suggests that Hat1 also has the potential to influence replication fork function. Studies in a wide range of eukaryotes show that loss of Hat1 sensitizes cells to DNA double strand breaks and causes HU sensitivity and genome instability in mammalian cells (19,57,58). In addition, it was recently reported that Hat1 is transiently recruited to chromatin during replication-coupled chromatin assembly and affects the protein composition of nascent chromatin(59). Therefore, we investigated whether Hat1 provides a link between the processing and assembly of newly synthesized histones and replication fork function. We confirm that Hat1 transiently associates with newly replicated DNA. We show that loss of Hat1 induces a dramatic reduction in replication fork progression and increases replication fork stalling. We also demonstrate that stalling of replication forks stabilizes the association of Hat1 with newly replicated DNA and that loss of Hat1 leads to destabilization of stalled forks and MRE11-dependent degradation of newly synthesized DNA.

## MATERIALS AND METHODS

### Cell culture conditions

Mouse embryonic fibroblasts were prepared as previously described (19). Cells were grown in DMEM (Sigma) supplemented with 10%FBS (Sigma) and Penicillin/Streptomycin (Gibco).

### Chromatin assembly assay

Three Hat1^+/+^ or Hat1^-/-^ MEF cell lines were seeded in equal quantities on coverslips and allowed to attach for 24 hours. For N-acetyl cysteine experiments (NAC), cells were seeded and allowed to grow for 48 hours in 5 mM NAC (Sigma; cat# A9165). Cells were then incubated with 10 uM IdU (Sigma; cat# I7125) for 30 minutes. For thymidine chases, IdU containing media was replaced with fresh media for indicated times. For cells treated with hydroxyurea (HU), IdU containing media was replaced with fresh media containing 4 mM HU (Sigma; cat# H8627) for designated times. 100 uM MIRIN (Sigma; cat# M9948) was added simultaneously where specified. Cells were then permeabilized with 0.5% TritonX-100 and fixed with 4% PFA simultaneously for 15 min, rinsed with PBS, and fixed again with 4% PFA for 10 minutes at room temperature. After several PBS washes, cells were incubated with 1N HCl for 10 minutes, washed with PBS until pH neutralizes, and blocked with 5% BSA for 1 hour at room temperature. BSA was removed with PBS washes and primary antibodies detecting IdU and a protein of interest were diluted in 1% BSA/0.3% TritonX-100 and added to cells overnight at 4°C (IdU: mouse anti-BrdU, 1:20, Becton Dickinson; Total Histone H4: rabbit anti-Total Histone H4, 1:500, Upstate Cell Signaling Solutions 05-858; Hat1: rabbit anti-Hat1, 1:1000, Abcam ab12163; H4K5Ac: rabbit anti-Histone H4 Lysine 5 Ac, 1:250, Abcam ab51997; H4K12Ac: rabbit anti-Histone H4 Lysine 12 Ac, 1:250, Abcam ab46983; PCNA: rabbit anti-PCNA, 1:50, Santa Cruz Biotechnology sc-7907; RAD51: rabbit anti-RAD51, 1:200, Abcam ab63801). The following day, primary antibodies were removed with PBS and cells were subjected to the Duolink™ Proximity Ligation Assay protocol according to manufacturer instructions (Sigma: DUO92008, DUO92004, DUO92002, DUO82049). After amplification, cells were incubated with AlexaFluor 488–conjugated anti-mouse secondary antibody (1:250, Molecular Probes) for 1 hour at room temperature, antibody was removed with PBS, nuclei were stained with 20 mM Hoechst 33342 Fluorescent Stain and mounted on slides using Vectashield. Slides were analyzed under an Andor Spinning Disk Confocal fluorescence microscope. Images were acquired using MetaMorph version 7.8.10 and quantification was completed using ImageJ version 1.52t according to a previously described protocol (63).

### DNA fiber assay

DNA was labeled with 50μM and 250μM for 20 minutes each. HU (Sigma) was used at 4mM for 5 hours; Mirin (Sigma) was used at 100μM for 5 hours. After labeling and treatment, cells were collected by trypsinization and resuspended in PBS. 2μL of the cells suspension were spotted on a glass slide and lysed with lysis buffer (0.5% SDS, 200 mM Tris-HCl, pH 7.4, 50 mM EDTA) for 10min, slides were then tilted to 15° to stretch the DNA fibers and fixed with Methanol/Acetic Acid (3:1) overnight at 4 degrees. Next day DNA was denatured with 2.5N HCl for 30min and wash several times with PBS before blocking with 1%BSA/PBS for 30min. Rat anti-BrdU (1:50, AbD Serotec) was used to detect CldU, and mouse anti-BrdU (1:20, Becton Dickinson) to detect IdU. Antibodies were diluted in blocking buffer and incubated for 1 hour at room temperature. AlexaFluor 594–conjugated anti-rat (1:250, Molecular Probes) and AlexaFluor 488–conjugated anti-mouse (1:250, Molecular Probes) were used as secondary antibodies and incubated for 1 hour at room temperature. Slides were mounted with Vectashield with DAPI.

### Immunofluorescence

Cells were seeded on coverslips and allowed to attach for 24 hours. Next day the cells were fixed with 4% PFA at room temperature for 10 minutes, washed several times with PBS and permeabilized with 0.5% Triton X-100/PBS for 15 minutes at room temperature, after several PBS washes cells were blocked with 5% BSA in PBS for 30 minutes at room temperature. Anti-phosphorylated ATR (Ser 428) (Cell Signaling #2853 1/100) was incubated overnight at 4 degrees. Next day after several washes, secondary AlexaFluor 594–conjugated anti-rabbit was diluted 1/250 and incubated 1 hour at room temperature. Antibody excess was extensively washed and slides were mounted with Vectashield with DAPI.

### Comet Assay

The Comet Assay kit (Trevigen, Gaitherburg,MD) was used according to the manufacture instructions. Briefly, MEFs were resuspended in ice cold PBS (Ca2+ and Mg2+ free) to a concentration of 1 × 10^5^ cells/ml. 5 µl cells were mixed with 50 µl of warm low melting Agarose and 50 µl were evenly spread onto the special comet slides. Slides were stored at 4 °C in the dark and transferred to pre-chilled lysis solution for 60 minutes at 4 °C. Next, slides were transferred to alkali unwinding solution at room temperature for 60 minutes. Slides were transferred to electrophoresis tank which contained pre-chilled Alkaline electrophoresis solution and run at 1 Volt/cm, 300 mA for 45 minutes at 4 degrees. The slides were immersed twice in deionized water for 5 minutes intervals and washed in 70% ethanol for 5 minutes. Then cells were stained with 100 µl of SYBR Green I for 5 minutes in the dark and slides were analyzed under Zeiss Axiophot fluorescence microscope. Images were taken using Metavue software version 6.3r2 software and comet tails were analyzed using opencomet by Imagej.

## RESULTS

### Hat1 transiently localizes to newly replicated DNA

Current models of replication-coupled chromatin assembly predict that Hat1 associates with, and modifies, newly synthesized histone H4 in the cytoplasm before transferring the modified histones to Asf1 for subsequent nuclear import and deposition. However, recent results using iPOND (isolation of <u>p</u>roteins <u>o</u>n <u>n</u>ascent <u>D</u>NA) suggested that Hat1 becomes transiently associated with newly replicated DNA(59). As this has the potential to significantly expand the role of Hat1 in genome duplication, we sought to confirm this observation.

Proximity ligation assay-based chromatin assembly assays (CAAs) have recently been developed and serve as a powerful method for analyzing protein dynamics on newly replicated DNA(60-63). The proximity ligation technique determines whether two molecules reside close to each other in the cell by employing two species-specific secondary antibodies that are fused to oligonucleotides. If the secondary antibodies recognize primary antibodies that are in close proximity, the oligonucleotides can both bind to a nicked circular DNA, creating a template for rolling circle replication. This amplifies sequences that can be bound by a fluorescent probe and visualized. To adapt this for use as a chromatin assembly assay, newly replicated DNA is labeled by the incorporation of the thymidine analog IdU. The proximity of proteins to newly replicated DNA is detected using antibodies against the protein of interest and antibodies recognizing IdU. To validate the CAA, we monitored the localization of PCNA, H4 lysine 5 acetylation and H4 lysine 12 acetylation to newly replicated DNA in Hat1^+/+^ and Hat1^-/-^ MEFs (mouse embryonic fibroblasts). As seen in Figure 1A, quantitation of the CAA precisely mirrored the results previously obtained with iPOND. The localization of PCNA to newly replicated DNA was Hat1-independent and the acetylation of H4 lysines 5 and 12 required Hat1(19,59).

**Figure 1.**
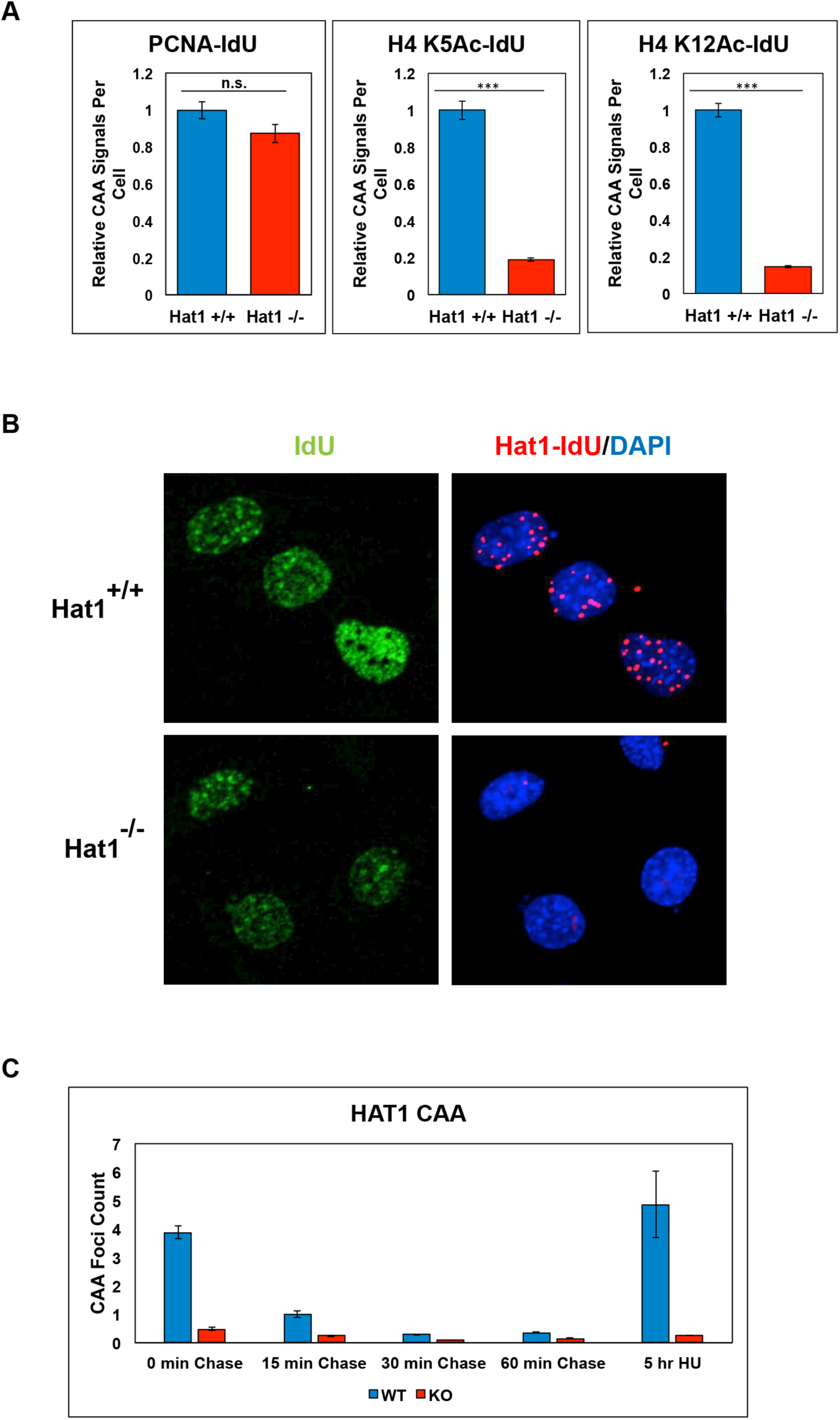
Hat1 localizes to newly replicated DNA. A) Quantitation of CAAs for PCNA, H4 lysine 5 acetylation and H4 lysine 12 acetylation performed in Hat1^+/+^ and Hat1^-/-^ cells (*** indicates p value < 0.01). B) Images of CAAs for Hat1 in Hat1^+/+^ and Hat1^-/-^ cells. Cells were visualized for IdU, Hat1 CAA and DAPI as indicated. C) Quantitation of Hat1 CAAs following a thymidine chase of the indicated length.

Using α-Hat1 antibodies, we tested whether Hat1 is in proximity to newly replicated DNA. There is abundant CAA signal in Hat1^+/+^ cells and only background in the Hat1^-/-^ cells (Figure 1B). We next asked whether Hat1 is transiently associated with newly synthesized DNA or whether it is stably bound to chromatin. We performed CAA assays immediately following a pulse of IdU and after 15, 30 and 60 minutes of a thymidine chase. As seen in Figure 1C, the level of Hat1 on newly replicated DNA is significantly reduced after a 15 minute chase and is completely lost after 30 minutes. Intriguingly, if replication forks are stalled by the addition of HU, Hat1 association with newly replicated DNA is stabilized for extended periods of time (at least 5 hours). These data verify that Hat1 is transiently associated with nascent chromatin near sites of DNA replication and becomes stably associated when replication forks stall.

### Hat1 is required for normal replication fork progression

The physical association of Hat1 with the highly dynamic chromatin at sites of DNA replication greatly expands the spectrum of potential functions for this enzyme in genome duplication. In particular, this raises the possibility that Hat1 plays a direct role in replication fork function or stability. To test this, we used DNA fiber analysis in Hat1^+/+^ and Hat1^-/-^ MEFs (Fig. 2*A*). Hat1^+/+^ and Hat1^-/-^ cells were incubated with CldU, followed by IdU incubation for equal times and replication fork progression was measured by DNA fiber analysis in which antibodies targeting the CldU (red) and IdU (green) are used to label the newly replicated DNA with different colors. The relative rates of replication fork progression were determined by measuring the lengths of the IdU tracts that are located at junctions with CldU labeled DNA, as this ensures that the replication fork was functional at the beginning of the IdU incubation. We observed a significant decrease in the length of labeled DNA fibers in the Hat1^-/-^ cells, indicating that DNA replication progressed more slowly in the absence of Hat1. Consistent with an effect of Hat1 loss on replication fork function, analysis of PCNA dynamics at the replication fork by CAA showed that PCNA dissociation is significantly delayed in the absence of Hat1 (Figure 2B).

**Figure 2.**
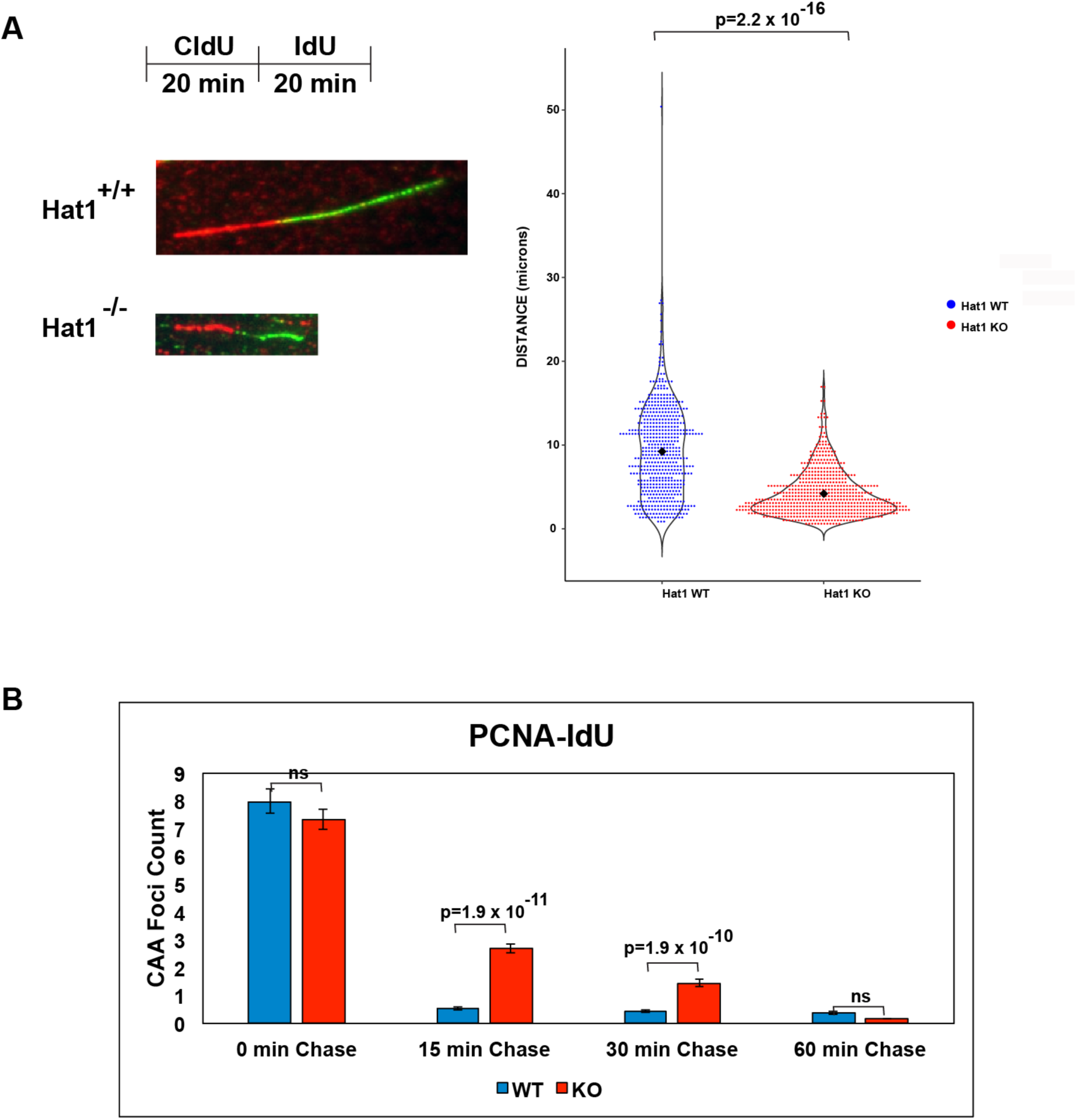
Hat1 is required for replication fork progression. A) Left top, schematic diagram of CldU/IdU labeling of DNA fibers. Left bottom, representative images of replication forks from Hat1^+/+^ and Hat1^-/-^ MEFs. Right, violin plot of the quantification of the fork distances traveled during CldU pulse (green fiber). At least 250 fibers were scored per cell line, P-value was calculated using the Wilcoxon test. B) Quantitation of PCNA CAAs following a thymidine chase for the indicated lengths of time.

### Loss of Hat1 increases replication fork stalling

A decreased rate of DNA replication can be due to decreases in the velocity of the replication fork or increases in the frequency of replication fork stalling. To test the latter possibility, we stained Hat1^+/+^ and Hat1^-/-^ cells with antibodies against phosphorylated ATR (Ser428). ATR is recruited to single stranded DNA at sites of replication fork stalling where it is activated by phosphorylation. Loss of Hat1 resulted in an increased number of cells positive for phospho-ATR foci (Fig. 3A).

**Figure 3.**
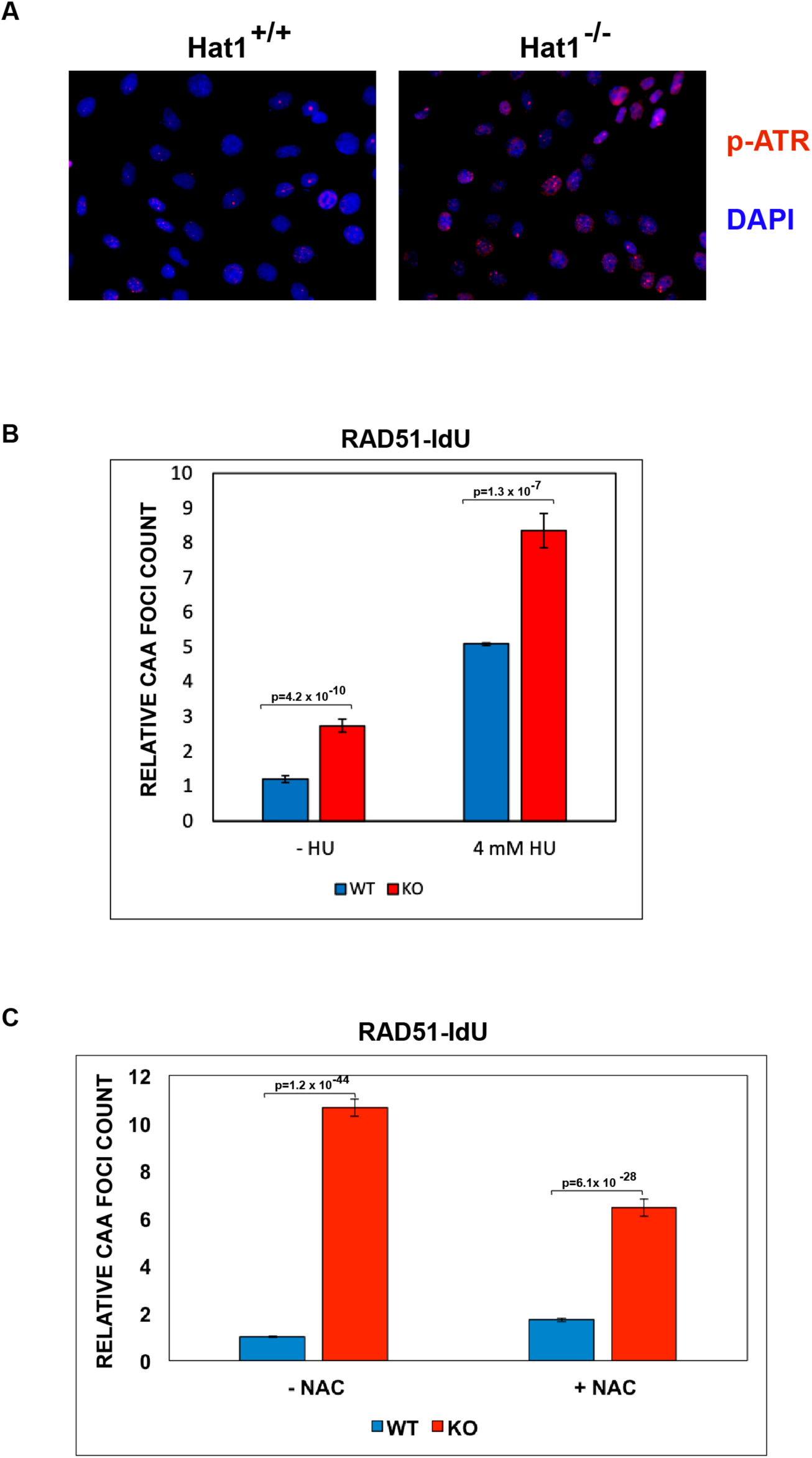
Hat1 loss causes replication fork stalling. A) phospho-ATR immunofluorescence (red) in Hat1^+/+^ and Hat1^-/-^ MEFs. Cells were also stained with DAPI. B) Quantitation of CAAs measuring localization of Rad51 to newly replicated DNA at stalled replication forks in Hat1^+/+^ and Hat1^-/-^ MEFs. C) Quantitation of Rad51 CAAs in Hat1+/+ and Hat1-/- cells grown in the absence and presence of 5 mM N-acetylcysteine (NAC).

To confirm the increase in replication fork stalling, we used the CAA to measure the association of Rad51 with the single strand DNA that is created at stalled replication forks(64). As seen in Figure 2B, there was a significant increase in the association of Rad51 with newly synthesized DNA in the absence of Hat1. As a positive control, treatment with HU results in a large increase in Rad51 CAA signal in the in both Hat1^+/+^ and Hat1^-/-^ cells.

The effect of Hat1 loss on replication fork stalling could be the result of increased levels of DNA damage inhibiting replication fork progression. Hat1^-/-^ MEFs contain increased levels of γ-H2AX, a marker of DNA damage(19). It was recently shown that Hat1^-/-^ MEFs display defects in mitochondrial function that result in elevated levels of ROS (reactive oxygen species). The mitochondrial defects are linked to the elevated levels of DNA damage as growth of Hat1^-/-^ MEFs in the presence of N-acetylcysteine (NAC) eliminated the elevated levels γ-H2AX staining(65). To determine whether Hat1 loss indirectly affects replication fork stalling through increased ROS and DNA damage, we performed Rad51 CAAs in Hat1^+/+^ and Hat1^-/-^ cells grown in the presence and absence of NAC. As seen in Figure 3C, growth in NAC led to a partial decrease in the Rad51 signal in Hat1^-/-^ cells. This suggests that loss of Hat1 effects replication fork stalling through both DNA damage-dependent and DNA damage-independent mechanisms. Together, these data indicate that Hat1 is necessary for proper replication fork function and the prevention of replication stress.

### Hat1 is critical for the stability of stalled replication forks

As seen in Figure 1C, Hat1 is stably associated with stalled replication forks. To determine whether Hat1 is involved in maintaining the stability of stalled replication forks, we analyzed the stability of newly replicated DNA at stalled replication forks using the DNA fiber assay. Hat1^+/+^ and Hat1^-/-^ cells were treated with CldU and IdU sequentially for equal lengths of time. HU was then added to induce replication fork stalling. After 5 hours, the lengths of the IdU and CldU tracts were measured. If the stalled replication forks remain stable, the ratio of IdU tract length to CldU tract length will be 1. If the newly replicated DNA (represented by the IdU labeled DNA) at the stalled forks is unstable, the ratio of IdU tract length to CldU tract length will be less than 1. As seen in Figure 4A, there was a significant decrease in the IdU tract length in the absence of Hat1. We conclude that Hat1 is required for the protection of newly replicated DNA at stalled replication forks.

**Figure 4.**
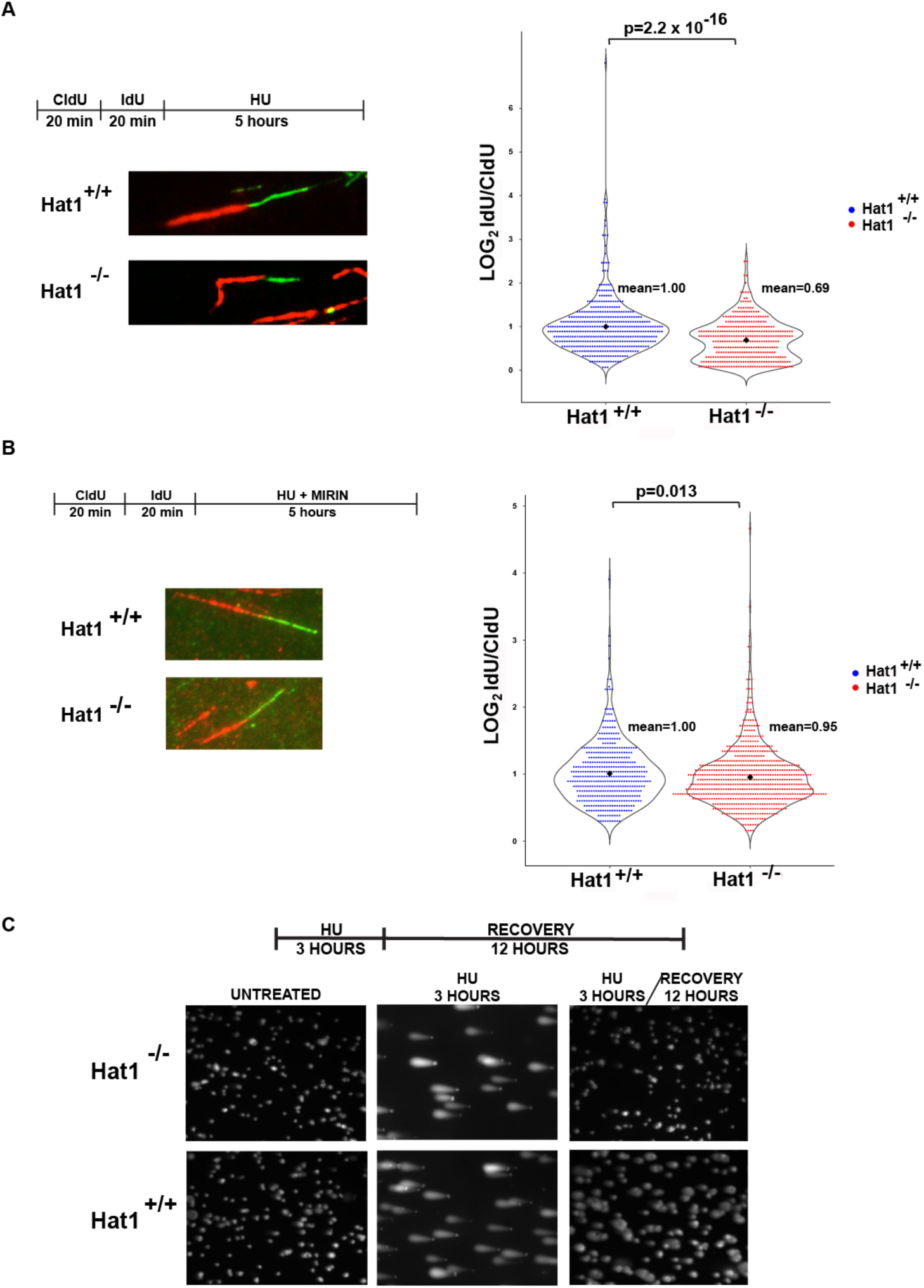
Hat1 protects newly replicated DNA from Mre11 digestion at stalled forks. A) Left top, schematic of CldU/IdU pulse-labeling followed by HU treatment. Left bottom, representative images of CldU and Idu replication forks in HU treated cells. Right, violin plot of IdU to CldU fiber length ratios for individual replication forks in HU treated Hat1^+/+^ and Hat1^-/-^ MEFs. At least 250 fibers were scored per cell line, P-value was calculated using the Wilcoxon test. B) Cells were treated as in (A) in the presence of 100μM Mirin. Left, representative pictures of both Hat1^+/+^ and Hat1^-/-^ DNA fibers. Right, violin plot of IdU to CldU fiber length ratios for individual replication forks. Statistical analysis was performed as in (A). C) Comet assay in Hat1^+/+^ and Hat1^-/-^ MEFs untreated or treated with HU and then released into fresh media for recovery.

The degradation of newly replicated DNA at stalled replication forks is the result of Mre11 nuclease activity (66)((67). To determine whether the instability of nascent DNA in the absence of Hat1 is also Mre11-dependent, Hat1^+/+^ and Hat1^-/-^ cells were sequentially treated with CldU and IdU for equal lengths of time. The cells were then treated with HU in the presence of Mirin, a specific inhibitor of Mre11 activity. As seen in Figure 4B, newly replicated DNA is equally stable in Hat1^+/+^ and Hat1^-/-^ cells when Mre11 activity is inhibited. These data indicate that Hat1 functions to protect newly replicated DNA from Mre11-mediated degradation.

We used a comet assay to determine whether Hat1-dependent replication fork instability led to a decrease in the ability of cells to recover from replication stress. Hat1^+/+^ and Hat1^-/-^ cells were treated with HU for 3 hours and then allowed to recover for 12 hours in the absence of HU. As seen in Figure 4C, Hat1^+/+^ cells were better able to recover from prolonged replication stress than the knock out cells, consistent with a loss of replication fork integrity in the absence of Hat1.

## DISCUSSION

Contrary to the predictions of current models of replication-coupled chromatin assembly, our results demonstrate that Hat1 localizes to chromatin at sites of DNA replication. There are several models to explain the localization of Hat1 to nascent chromatin. First, Hat1 may not transfer H3/H4 dimers to Asf1. Rather, Hat1 may remain associated with the H3/H4 dimers throughout the entire replication-coupled chromatin assembly process and load onto newly replicated DNA through CAF-1-mediated deposition of H3/H4/Hat1 complexes. This model is consistent with numerous proteomic studies that have identified the Hat1 complex as major components of soluble H3 and H4 complexes(22,26,68-72). Alternatively, following transfer of H3/H4 dimers to Asf1, Hat1 may enter the nucleus independently and bind to nascent chromatin after histone deposition. Finally, Hat1 may participate in an additional chromatin assembly pathway distinct from the Asf1/CAF-1 pathway. One potential pathway may utilize the histone chaperone NASP. A distinct nuclear yeast Hat1 complex contains histones H3 and H4 and a histone chaperone, Hif1, which is the yeast homolog of NASP(15,21,68). Subsequent experiments have shown that NASP also interacts with the Hat1 complex in mammalian cells(22). NASP is important for buffering the pools of soluble H3/H4, particularly under conditions of replication stress, and can form a multi-chaperone complex with Asf1. Several studies have shown that NASP can function as a nucleosome assembly factor *in vitro*(73-75). Hence, this model predicts that Hat1 localizes to newly replicated DNA in conjunction with NASP-mediated deposition of H3/H4.

It is clear that DNA replication is coupled to newly synthesized histone deposition. This link was originally suggested by studies demonstrating that DNA replication required active protein synthesis(76-78). More recently, histone supply was more directly linked to replication fork function by experiments that specifically limited histone production(48,50). The assembly of newly synthesized histones into chromatin was directly implicated in replication fork function through the identification of DNA replication defects in cells lacking components of the replication-coupled chromatin assembly pathway, such as Asf1 and CAF-1(39,49,51).

While histone deposition in the wake of the replication fork is clearly important for fork progression, histone deposition occurs with normal kinetics in Hat1^-/-^ cells, leaving open the question of how Hat1 influences replication fork progression(59). There are a number of possibilities. While histones are still deposited onto newly replicated DNA in the absence of Hat1, nucleosomes may not be assembled properly. Proteomic analysis showed that nascent chromatin assembled in the absence of Hat1 was associated with increased levels of topoisomerase 2 and Hat1^-/-^ cells are hypersensitive to topoisomerase 2 inhibition(59). Therefore, topological defects in nascent chromatin assembled in the absence of Hat1 may impede replication fork progression. The nascent chromatin proteomics also identified a number of proteins that are depleted from nascent chromatin assembled in Hat1^-/-^ cells. These include the bromodomain proteins, Baz1a, Brg1 and Brd3(59). Decreased levels of these proteins in the proximity of replication forks may create an altered chromatin structure that negatively affects replication fork function or they may be directly involved in replisome function. Indeed, it was recently shown that Brd2, Brd3 and Brd4 function at the replication fork to antagonize the ATAD5-mediated unloading of PCNA, which is consistent with our observation that PCNA unloading is slowed in Hat1^-/-^ cells(79). Finally, the presence of Hat1 on nascent chromatin near replication forks suggests that Hat1 may directly modify and regulate components of the replisome.

The association of Hat1 with nascent chromatin is transient but becomes stable if replication forks stall. The physical association of Hat1 with stalled forks is likely to be functionally relevant as Hat1 is required for the stability of stalled replication forks. An attractive mechanism for the role of Hat1 in replication fork stabilization involves the recruitment of Rad51 to stalled forks. Rad51 binds to single strand DNA at stalled replication forks and plays a central role in maintaining replication fork stability. Hat1 forms an S-phase-specific complex with Rad51 and is involved in the recruitment of Rad51 to DNA double strand breaks (80). However, we do not detect any decrease in Rad51 localization to stalled replication forks in Hat1^-/-^ cells, suggesting that the mechanisms for Rad51 recruitment to DNA double strand breaks and stalled replication forks are distinct.

It has also been suggested that Hat1 is involved in the initiation of DNA replication. Studies in yeast showed that Hat1 physically interacts with the origin recognition complex (ORC). In addition, combining mutations in Hat1 with temperature sensitive alleles of ORC components or CDC45 resulted in synthetic growth defects. Hat1 was also recruited to origins of replication at the time of origin activation. Despite these connections, there were no defects in replication origin firing in Hat1 mutants in yeast(81).

Our results suggest an update to current models of replication-coupled chromatin assembly to incorporate the localization of Hat1 to nascent chromatin at sites of DNA replication. In addition, our results indicate that Hat1 lays a direct and integral role in both genome and epigenome duplication.

## Acknowledgements

This work was support by a grant form the National Institutes of Health (R01 GM062970 to M.R.P.)). Microscopy was supported by a grant from the NIH/NINDS (P30 NS104177).

## References

1. Franco, A. A., Lam, W. M., Burgers, P. M., and Kaufman, P. D. (2005) Histone deposition protein Asf1 maintains DNA replisome integrity and interacts with replication factor C. Genes Dev 19, 1365–1375

2. Clement, C., and Almouzni, G. (2015) MCM2 binding to histones H3-H4 and ASF1 supports a tetramer-to-dimer model for histone inheritance at the replication fork. Nat Struct Mol Biol 22, 587–589

3. Richet, N., Liu, D., Legrand, P., Velours, C., Corpet, A., Gaubert, A., Bakail, M., Moal-Raisin, G., Guerois, R., Compper, C., Besle, A., Guichard, B., Almouzni, G., and Ochsenbein, F. (2015) Structural insight into how the human helicase subunit MCM2 may act as a histone chaperone together with ASF1 at the replication fork. Nucleic Acids Res 43, 1905–1917

4. Huang, H., Stromme, C. B., Saredi, G., Hodl, M., Strandsby, A., Gonzalez-Aguilera, C., Chen, S., Groth, A., and Patel, D. J. (2015) A unique binding mode enables MCM2 to chaperone histones H3-H4 at replication forks. Nat Struct Mol Biol 22, 618–626

5. Wang, H., Wang, M., Yang, N., and Xu, R. M. (2015) Structure of the quaternary complex of histone H3-H4 heterodimer with chaperone ASF1 and the replicative helicase subunit MCM2. Protein & cell 6, 693–697

6. Liu, S., Xu, Z., Leng, H., Zheng, P., Yang, J., Chen, K., Feng, J., and Li, Q. (2017) RPA binds histone H3-H4 and functions in DNA replication-coupled nucleosome assembly. Science 355, 415–420

7. Jasencakova, Z., Scharf, A. N., Ask, K., Corpet, A., Imhof, A., Almouzni, G., and Groth, A. (2010) Replication stress interferes with histone recycling and predeposition marking of new histones. Mol Cell 37, 736–743

8. Jasencakova, Z., and Groth, A. (2010) Restoring chromatin after replication: how new and old histone marks come together. Semin Cell Dev Biol 21, 231–237

9. Groth, A., Corpet, A., Cook, A. J., Roche, D., Bartek, J., Lukas, J., and Almouzni, G. (2007) Regulation of replication fork progression through histone supply and demand. Science 318, 1928–1931

10. Bellelli, R., Belan, O., Pye, V. E., Clement, C., Maslen, S. L., Skehel, J. M., Cherepanov, P., Almouzni, G., and Boulton, S. J. (2018) POLE3-POLE4 Is a Histone H3-H4 Chaperone that Maintains Chromatin Integrity during DNA Replication. Mol Cell 72, 112–126 e115

11. Foltman, M., Evrin, C., De Piccoli, G., Jones, R. C., Edmondson, R. D., Katou, Y., Nakato, R., Shirahige, K., and Labib, K. (2013) Eukaryotic replisome components cooperate to process histones during chromosome replication. Cell Rep 3, 892–904

12. Yang, J., Zhang, X., Feng, J., Leng, H., Li, S., Xiao, J., Liu, S., Xu, Z., Xu, J., Li, D., Wang, Z., Wang, J., and Li, Q. (2016) The Histone Chaperone FACT Contributes to DNA Replication-Coupled Nucleosome Assembly. Cell Rep 16, 3414

13. Annunziato, A. T. (2012) Assembling chromatin: The long and winding road. Biochim Biophys Acta 1819, 196–210

14. Rivera, C., Saavedra, F., Alvarez, F., Díaz-Celis, C., Ugalde, V., Li, J., Forné, I., Gurard-Levin, Z. A., Almouzni, G., Imhof, A., and Loyola, A. (2015) Methylation of histone H3 lysine 9 occurs during translation. Nucleic Acids Research 43, 9097–9106

15. Campos, E. I., Fillingham, J., Li, G., Zheng, H., Voigt, P., Kuo, W. H., Seepany, H., Gao, Z., Day, L. A., Greenblatt, J. F., and Reinberg, D. The program for processing newly synthesized histones H3.1 and H4. Nat Struct Mol Biol 17, 1343–1351

16. Parthun, M. R., Widom, J., and Gottschling, D. E. (1996) The major cytoplasmic histone acetyltransferase in yeast: links to chromatin replication and histone metabolism. Cell 87, 85–94

17. Kleff, S., Andrulis, E. D., Anderson, C. W., and Sternglanz, R. (1995) Identification of a gene encoding a yeast histone H4 acetyltransferase. J Biol Chem 270, 24674–24677

18. Verreault, A., Kaufman, P. D., Kobayashi, R., and Stillman, B. (1998) Nucleosomal DNA regulates the core-histone-binding subunit of the human Hat1 acetyltransferase. Curr Biol 8, 96–108

19. Nagarajan, P., Ge, Z., Sirbu, B., Doughty, C., Agudelo Garcia, P. A., Schlederer, M., Annunziato, A. T., Cortez, D., Kenner, L., and Parthun, M. R. (2013) Histone acetyl transferase 1 is essential for mammalian development, genome stability, and the processing of newly synthesized histones H3 and H4. PLoS Genet 9, e1003518

20. Chang, L., Loranger, S. S., Mizzen, C., Ernst, S. G., Allis, C. D., and Annunziato, A. T. (1997) Histones in transit: cytosolic histone complexes and diacetylation of H4 during nucleosome assembly in human cells. Biochemistry 36, 469–480

21. Fillingham, J., Recht, J., Silva, A. C., Suter, B., Emili, A., Stagljar, I., Krogan, N. J., Allis, C. D., Keogh, M. C., and Greenblatt, J. F. (2008) Chaperone control of the activity and specificity of the histone H3 acetyltransferase Rtt109. Mol Cell Biol 28, 4342–4353

22. Campos, E. I., Fillingham, J., Li, G., Zheng, H., Voigt, P., Kuo, W. H., Seepany, H., Gao, Z., Day, L. A., Greenblatt, J. F., and Reinberg, D. (2010) The program for processing newly synthesized histones H3.1 and H4. Nat Struct Mol Biol 17, 1343–1351

23. Alvarez, F., Munoz, F., Schilcher, P., Imhof, A., Almozuni, G., and Loyola, A. (2011) Sequential establishment of marks on soluble histones H3 and H4. J Biol Chem

24. Blackwell, J. S., Jr., Wilkinson, S. T., Mosammaparast, N., and Pemberton, L. F. (2007) Mutational analysis of H3 and H4 N termini reveals distinct roles in nuclear import. J Biol Chem 282, 20142–20150

25. Mosammaparast, N., Ewart, C. S., and Pemberton, L. F. (2002) A role for nucleosome assembly protein 1 in the nuclear transport of histones H2A and H2B. Embo J 21, 6527–6538

26. Barman, H. K., Takami, Y., Nishijima, H., Shibahara, K., Sanematsu, F., and Nakayama, T. (2008) Histone acetyltransferase-1 regulates integrity of cytosolic histone H3-H4 containing complex. Biochem Biophys Res Commun 373, 624–630

27. Adkins, M. W., Carson, J. J., English, C. M., Ramey, C. J., and Tyler, J. K. (2007) The histone chaperone anti-silencing function 1 stimulates the acetylation of newly synthesized histone H3 in S-phase. J Biol Chem 282, 1334–1340

28. Burgess, R. J., Zhou, H., Han, J., and Zhang, Z. (2010) A role for Gcn5 in replication-coupled nucleosome assembly. Mol Cell 37, 469–480

29. Chen, C. C., Carson, J. J., Feser, J., Tamburini, B., Zabaronick, S., Linger, J., and Tyler, J. K. (2008) Acetylated lysine 56 on histone H3 drives chromatin assembly after repair and signals for the completion of repair. Cell 134, 231–243

30. Ge, Z., Nair, D., Guan, X., Rastogi, N., Freitas, M. A., and Parthun, M. R. (2013) Sites of acetylation on newly synthesized histone h4 are required for chromatin assembly and DNA damage response signaling. Mol Cell Biol 33, 3286–3298

31. Han, J., Zhou, H., Horazdovsky, B., Zhang, K., Xu, R. M., and Zhang, Z. (2007) Rtt109 acetylates histone H3 lysine 56 and functions in DNA replication. Science 315, 653–655

32. Han, J., Zhou, H., Li, Z., Xu, R. M., and Zhang, Z. (2007) Acetylation of lysine 56 of histone H3 catalyzed by RTT109 and regulated by ASF1 is required for replisome integrity. J Biol Chem 282, 28587–28596

33. Han, J., Zhou, H., Li, Z., Xu, R. M., and Zhang, Z. (2007) The Rtt109-Vps75 histone acetyltransferase complex acetylates non-nucleosomal histone H3. J Biol Chem 282, 14158–14164

34. Kaplan, T., Liu, C. L., Erkmann, J. A., Holik, J., Grunstein, M., Kaufman, P. D., Friedman, N., and Rando, O. J. (2008) Cell cycle- and chaperone-mediated regulation of H3K56ac incorporation in yeast. PLoS Genet 4, e1000270

35. Recht, J., Tsubota, T., Tanny, J. C., Diaz, R. L., Berger, J. M., Zhang, X., Garcia, B. A., Shabanowitz, J., Burlingame, A. L., Hunt, D. F., Kaufman, P. D., and Allis, C. D. (2006) Histone chaperone Asf1 is required for histone H3 lysine 56 acetylation, a modification associated with S phase in mitosis and meiosis. Proc Natl Acad Sci U S A 103, 6988–6993

36. Tsubota, T., Berndsen, C. E., Erkmann, J. A., Smith, C. L., Yang, L., Freitas, M. A., Denu, J. M., and Kaufman, P. D. (2007) Histone H3-K56 acetylation is catalyzed by histone chaperone-dependent complexes. Mol Cell 25, 703–712

37. Das, C., Lucia, M. S., Hansen, K. C., and Tyler, J. K. (2009) CBP/p300-mediated acetylation of histone H3 on lysine 56. Nature 459, 113–117

38. Li, Q., Zhou, H., Wurtele, H., Davies, B., Horazdovsky, B., Verreault, A., and Zhang, Z. (2008) Acetylation of histone H3 lysine 56 regulates replication-coupled nucleosome assembly. Cell 134, 244–255

39. Hoek, M., and Stillman, B. (2003) Chromatin assembly factor 1 is essential and couples chromatin assembly to DNA replication in vivo. Proc Natl Acad Sci U S A 100, 12183–12188

40. Malay, A. D., Umehara, T., Matsubara-Malay, K., Padmanabhan, B., and Yokoyama, S. (2008) Crystal structures of fission yeast histone chaperone Asf1 complexed with the Hip1 B-domain or the Cac2 C terminus. J Biol Chem 283, 14022–14031

41. Mello, J. A., Sillje, H. H., Roche, D. M., Kirschner, D. B., Nigg, E. A., and Almouzni, G. (2002) Human Asf1 and CAF-1 interact and synergize in a repair-coupled nucleosome assembly pathway. EMBO Rep 3, 329–334

42. Moggs, J. G., Grandi, P., Quivy, J. P., Jonsson, Z. O., Hubscher, U., Becker, P. B., and Almouzni, G. (2000) A CAF-1-PCNA-mediated chromatin assembly pathway triggered by sensing DNA damage. Mol Cell Biol 20, 1206–1218.

43. Ridgway, P., and Almouzni, G. (2000) CAF-1 and the inheritance of chromatin states: at the crossroads of DNA replication and repair. J Cell Sci 113, 2647–2658.

44. Shibahara, K., and Stillman, B. (1999) Replication-dependent marking of DNA by PCNA facilitates CAF-1-coupled inheritance of chromatin. Cell 96, 575–585

45. Tyler, J. K., Collins, K. A., Prasad-Sinha, J., Amiott, E., Bulger, M., Harte, P. J., Kobayashi, R., and Kadonaga, J. T. (2001) Interaction between the Drosophila CAF-1 and ASF1 chromatin assembly factors. Mol Cell Biol 21, 6574–6584

46. Liu, W. H., Roemer, S. C., Port, A. M., and Churchill, M. E. (2012) CAF-1-induced oligomerization of histones H3/H4 and mutually exclusive interactions with Asf1 guide H3/H4 transitions among histone chaperones and DNA. Nucleic Acids Res 40, 11229–11239

47. Winkler, D. D., Zhou, H., Dar, M. A., Zhang, Z., and Luger, K. (2012) Yeast CAF-1 assembles histone (H3-H4)2 tetramers prior to DNA deposition. Nucleic Acids Res 40, 10139–10149

48. Mejlvang J. F. Y., Alabert, C., Neelsen K.J., Jasencakova Z, Zhao, X.,Lees M., Sandelin A., Pasero P., Lopes M. and Groth A. (2014) New Histone Supply Regulates Replication Fork Speek and PCNA Loading. Journal of Cell Biology 204, 29–43

49. Clemente-Ruiz, M., Gonzalez-Prieto, R., and Prado, F. (2011) Histone H3K56 acetylation, CAF1, and Rtt106 coordinate nucleosome assembly and stability of advancing replication forks. PLoS Genet 7, e1002376

50. Clemente-Ruiz, M., and Prado, F. (2009) Chromatin assembly controls replication fork stability. EMBO Rep 10, 790–796

51. Takami, Y., Ono, T., Fukagawa, T., Shibahara, K., and Nakayama, T. (2007) Essential role of chromatin assembly factor-1-mediated rapid nucleosome assembly for DNA replication and cell division in vertebrate cells. Mol Biol Cell 18, 129–141

52. Bhaskara, S., Jacques, V., Rusche, J. R., Olson, E. N., Cairns, B. R., and Chandrasekharan, M. B. (2013) Histone deacetylases 1 and 2 maintain S-phase chromatin and DNA replication fork progression. Epigenetics Chromatin 6, 27

53. Kehrli, K., Phelps, M., Lazarchuk, P., Chen, E., Monnat, R., Jr., and Sidorova, J. M. (2016) Class I Histone Deacetylase HDAC1 and WRN RECQ Helicase Contribute Additively to Protect Replication Forks upon Hydroxyurea-induced Arrest. J Biol Chem 291, 24487–24503

54. Sirbu, B. M., Couch, F. B., Feigerle, J. T., Bhaskara, S., Hiebert, S. W., and Cortez, D. (2011) Analysis of protein dynamics at active, stalled, and collapsed replication forks. Genes Dev 25, 1320–1327

55. Milutinovic, S., Zhuang, Q., and Szyf, M. (2002) Proliferating cell nuclear antigen associates with histone deacetylase activity, integrating DNA replication and chromatin modification. J Biol Chem 277, 20974–20978

56. Lazarchuk, P., Hernandez-Villanueva, J., Pavlova, M. N., Federation, A., MacCoss, M., and Sidorova, J. M. (2020) Mutual balance of histone deacetylases HDAC1, HDAC2, and the acetyl reader ATAD2 regulates the level of acetylation of histone H4 on nascent chromatin of human cells. Mol Cell Biol

57. Qin, S., and Parthun, M. R. (2006) Recruitment of the type B histone acetyltransferase Hat1p to chromatin is linked to DNA double-strand breaks. Mol Cell Biol 26, 3649–3658

58. Qin, S., and Parthun, M. R. (2002) Histone H3 and the histone acetyltransferase Hat1p contribute to DNA double-strand break repair. Mol Cell Biol 22, 8353–8365

59. Agudelo Garcia, P. A., Hoover, M. E., Zhang, P., Nagarajan, P., Freitas, M. A., and Parthun, M. R. (2017) Identification of multiple roles for histone acetyltransferase 1 in replication-coupled chromatin assembly. Nucleic Acids Res 45, 9319–9335

60. Roy, S., Luzwick, J. W., and Schlacher, K. (2018) SIRF: Quantitative in situ analysis of protein interactions at DNA replication forks. J Cell Biol 217, 1521–1536

61. Petruk, S., Sedkov, Y., Johnston, D. M., Hodgson, J. W., Black, K. L., Kovermann, S. K., Beck, S., Canaani, E., Brock, H. W., and Mazo, A. (2012) TrxG and PcG proteins but not methylated histones remain associated with DNA through replication. Cell 150, 922–933

62. Petruk, S., Cai, J., Sussman, R., Sun, G., Kovermann, S. K., Mariani, S. A., Calabretta, B., McMahon, S. B., Brock, H. W., Iacovitti, L., and Mazo, A. (2017) Delayed Accumulation of H3K27me3 on Nascent DNA Is Essential for Recruitment of Transcription Factors at Early Stages of Stem Cell Differentiation. Mol Cell 66, 247–257 e245

63. Lazarchuk, P., Roy, S., Schlacher, K., and Sidorova, J. (2019) Detection and Quantitation of Acetylated Histones on Replicating DNA Using In Situ Proximity Ligation Assay and Click-It Chemistry. Methods Mol Biol 1983, 29–45

64. Bhat, K. P., and Cortez, D. (2018) RPA and RAD51: fork reversal, fork protection, and genome stability. Nat Struct Mol Biol 25, 446–453

65. Nagarajan, P., Agudelo Garcia, P. A., Iyer, C. C., Popova, L. V., Arnold, W. D., and Parthun, M. R. (2019) Early-onset aging and mitochondrial defects associated with loss of histone acetyltransferase 1 (Hat1). Aging cell, e12992

66. Hashimoto, Y., Ray Chaudhuri, A., Lopes, M., and Costanzo, V. (2010) Rad51 protects nascent DNA from Mre11-dependent degradation and promotes continuous DNA synthesis. Nat Struct Mol Biol 17, 1305–1311

67. Kolinjivadi, A. M., Sannino, V., De Antoni, A., Zadorozhny, K., Kilkenny, M., Técher, H., Baldi, G., Shen, R., Ciccia, A., Pellegrini, L., Krejci, L., and Costanzo, V. (2017) Smarcal1-Mediated Fork Reversal Triggers Mre11-Dependent Degradation of Nascent DNA in the Absence of Brca2 and Stable Rad51 Nucleofilaments. Mol Cell 67, 867-881.e867

68. Ai, X., and Parthun, M. R. (2004) The nuclear Hat1p/Hat2p complex: a molecular link between type B histone acetyltransferases and chromatin assembly. Mol Cell 14, 195–205

69. Ejlassi-Lassallette, A., Mocquard, E., Arnaud, M. C., and Thiriet, C. (2011) H4 replication-dependent diacetylation and Hat1 promote S-phase chromatin assembly in vivo. Mol Biol Cell 22, 245–255

70. Tagami, H., Ray-Gallet, D., Almouzni, G., and Nakatani, Y. (2004) Histone H3.1 and H3.3 complexes mediate nucleosome assembly pathways dependent or independent of DNA synthesis. Cell 116, 51–61

71. Drane, P., Ouararhni, K., Depaux, A., Shuaib, M., and Hamiche, A. (2010) The death-associated protein DAXX is a novel histone chaperone involved in the replication-independent deposition of H3.3. Genes Dev 24, 1253–1265

72. Saade, E., Mechold, U., Kulyyassov, A., Vertut, D., Lipinski, M., and Ogryzko, V. (2009) Analysis of interaction partners of H4 histone by a new proteomics approach. Proteomics 9, 4934–4943

73. Wang, H., Ge, Z., Walsh, S. T., and Parthun, M. R. (2012) The human histone chaperone sNASP interacts with linker and core histones through distinct mechanisms. Nucleic Acids Res 40, 660–669

74. Wang, H., Walsh, S. T., and Parthun, M. R. (2008) Expanded binding specificity of the human histone chaperone NASP. Nucleic Acids Res 36, 5763–5772

75. Finn, R. M., Browne, K., Hodgson, K. C., and Ausio, J. (2008) sNASP, a histone H1-specific eukaryotic chaperone dimer that facilitates chromatin assembly. Biophys J

76. Weintraub, H. (1972) A possible role for histone in the synthesis of DNA. Nature 240, 449–453

77. Weintraub, H., and Holtzer, H. (1972) Fine control of DNA synthesis in developing chick red blood cells. J Mol Biol 66, 13–35

78. Seale, R. L., and Simpson, R. T. (1975) Effects of cycloheximide on chromatin biosynthesis. J Mol Biol 94, 479–501

79. Wessel, S. R., Mohni, K. N., Luzwick, J. W., Dungrawala, H., and Cortez, D. (2019) Functional Analysis of the Replication Fork Proteome Identifies BET Proteins as PCNA Regulators. Cell Rep 28, 3497–3509 e3494

80. Yang, X., Li, L., Liang, J., Shi, L., Yang, J., Yi, X., Zhang, D., Han, X., Yu, N., and Shang, Y. (2013) Histone acetyltransferase 1 promotes homologous recombination in DNA repair by facilitating histone turnover. J Biol Chem 288, 18271–18282

81. Suter, B., Pogoutse, O., Guo, X., Krogan, N., Lewis, P., Greenblatt, J. F., Rine, J., and Emili, A. (2007) Association with the origin recognition complex suggests a novel role for histone acetyltransferase Hat1p/Hat2p. BMC Biol 5, 38

